# Activation of serotonin neurons promotes active exploitation in a probabilistic foraging task

**DOI:** 10.1101/170472

**Authors:** Eran Lottem, Dhruba Banerjee, Pietro Vertechi, Dario Sarra, Matthijs oude Lohuis, Zachary F. Mainen

## Abstract

The neuromodulator serotonin (5-HT) has been implicated in a variety of functions that involve patience or impulse control. For example, activation of 5-HT neurons promotes waiting for delayed rewards. Many of these effects are consistent with a long-standing theory that 5-HT promotes behavioral inhibition, a motivational bias favoring passive over active behaviors. To further test this idea, we studied the impact of 5-HT in a probabilistic foraging task, in which mice must learn the statistics of the environment and infer when to leave a depleted foraging site for the next. Critically, mice were required to actively nose poke in order to exploit a given site. We found that optogenetic activation of 5-HT neurons in the dorsal raphe nucleus increased the willingness of mice to actively attempt to exploit a reward site before giving up. These results indicate that behavioral inhibition is not an adequate description of 5-HT function and suggest that a unified account must be based on a higher-order function.

---

Serotonin (5-HT) is a central neuromodulator that is implicated in the regulation of many processes and is one of the most important targets of psychoactive drugs^1,2^. As a unifying concept for 5-HT’s manifold effects, Soubrié^3^ put forward the hypothesis that a major function of 5-HT is to promote behavioral inhibition. Building on work showing that blockade of serotonin transmission results in continued responses to stimuli that are no longer rewarding^4–7^, he argued that in situations wherein animals face a decision between active response and passivity, higher levels of 5-HT would bias the decision in favor of the latter.

More recently, the study of 5-HT and behavioral inhibition has concentrated chiefly on impulse control^8–10^. One of the most common tasks used to study impulsive behavior is the five-choice serial reaction time task (5-CSRTT), in which rodents are required to respond to visual stimuli to obtain rewards^11^. Although this task was not specifically designed to measure impulsivity, animals sometimes respond prematurely (i.e. before stimulus presentation), a behavior indicative of impulsivity. Consistent with the idea that 5-HT promotes patience, alterations such as brain-wide 5-HT depletion increase impulsivity in this task^12^.

Another line of experiments focuses on animals’ ability to wait in order to obtain reward^13^. Electrophysiological recordings from the dorsal raphe nucleus (DRN; the major source of serotonin to the forebrain) in rats trained to wait for delayed rewards have found that most of these neurons increase their firing rates during waiting^14^. Moreover, pharmacological blocking of these neurons by the local application of the 5-HT_1A_ receptor agonist 8-hydroxy-2-(di-n-propylamino) tetralin (8-OH-DPAT) locally into the DRN promotes premature leaving^14^, whereas optogenetic activation of the same neurons promotes patience^15,16^.

All of these results are still broadly consistent with the Soubrié theory of behavioral inhibition. By reducing the motivation to act, increased serotonin levels would reduce the rate of premature responses in the 5-CSRTT, and increase the time animals can wait to obtain delayed rewards. However, they are also consistent with a different theory, in which serotonin promotes not passivity or the ability to tolerate inaction, but the patience or persistence to carry out an action that is itself costly or unrewarding but helps lead to delayed benefit. Thus, successful waiting, in this alternative account, consists not only in suppressing the urge to respond prematurely, but actively carrying out a behavioral alternative to responding. Indeed, children facing the “marshmallow task”, in which they must keep from eating one tasty treat in order to gain a second one later on, often succeed not only by passively controlling their impulse to eat the available marshmallow, but by actively distracting themselves by performing alternative behaviors^17^. So far, the above-mentioned tasks used to study serotonin, waiting and passivity coincide by design and so these alternative explanations are not readily distinguishable.

Here, we sought to disambiguate whether serotonin promotes waiting through behavioral inhibition or though persistence in the context of foraging behavior. In natural foraging situations, resources are typically found in isolated patches that become exhausted over time, facing animals with an exploitation/exploration dilemma: they must choose between working within in a given patch and giving up to travel to a different one. When foraging within a patch is itself subject to uncertainty—that is the income from a patch is itself irregular or probabilistic—then exploitation requires patience. But this kind of patience is active rather than passive: in order to obtain rewards animals must continue to seek for food even when such attempts are sometimes unsuccessful.

We developed a probabilistic foraging task in which mice nose-poke at one of two nose pokes and are rewarded for their pokes (foraging attempts) according to a random probability schedule that decays exponentially to zero. Due to the probabilistic nature of the reward, in each trial, a mouse will experience rewarded pokes interspersed with unrewarded pokes. Each sequence of one or more omitted rewards challenges the mouse with a waiting task, but one that requires active behavior. We reasoned that it if 5-HT promotes behavioral inhibition and passive behavior, then it should suppress poking, resulting in fewer pokes at each port visit. If, instead, 5-HT promotes persistence, it should promote the active behavior, or poking, which is required to obtain rewards. Indeed, in the nematode worm *C. elegans* it was found that 5-HT favors dwelling in a food patch over roaming^18^.

We found that optogenetic activation of DRN 5-HT neurons enhanced the active exploitation of the current resource. It increased the number of active nose-pokes a mouse would carry out in an attempt to gain water before giving up. These results contradict the behavioral inhibition hypothesis and support the notion that 5-HT promotes waiting by enhancing persistence in the face of uncertainty and delay.

## Results

### Mouse behavior in a probabilistic foraging task

To study the role of 5-HT in foraging mice, we developed a novel probabilistic foraging task (PFT). Water-restricted mice were placed in an elongated rectangular chamber containing two nose poke water ports, one at each end (Fig. 1a), which represented foraging sites. Body position was determined using video tracking. We defined a region of interest (ROI) around each port. A foraging “trial” was considered as a visit to one port, defined as the period between ROI entry and exit (Fig. 1b,c). Correct trials were ones in which the mouse alternated between ports, whereas error trials were those in which the mouse left and then reentered the same ROI with no visit to the other port in between. In correct trials, each nose poke into the port was either rewarded or not according to a defined probability schedule (Fig. 1d). Reward probabilities were reset to their highest value at the start of each trial and declined exponentially with each poke. Error trials were unrewarded.

**Figure 1.**
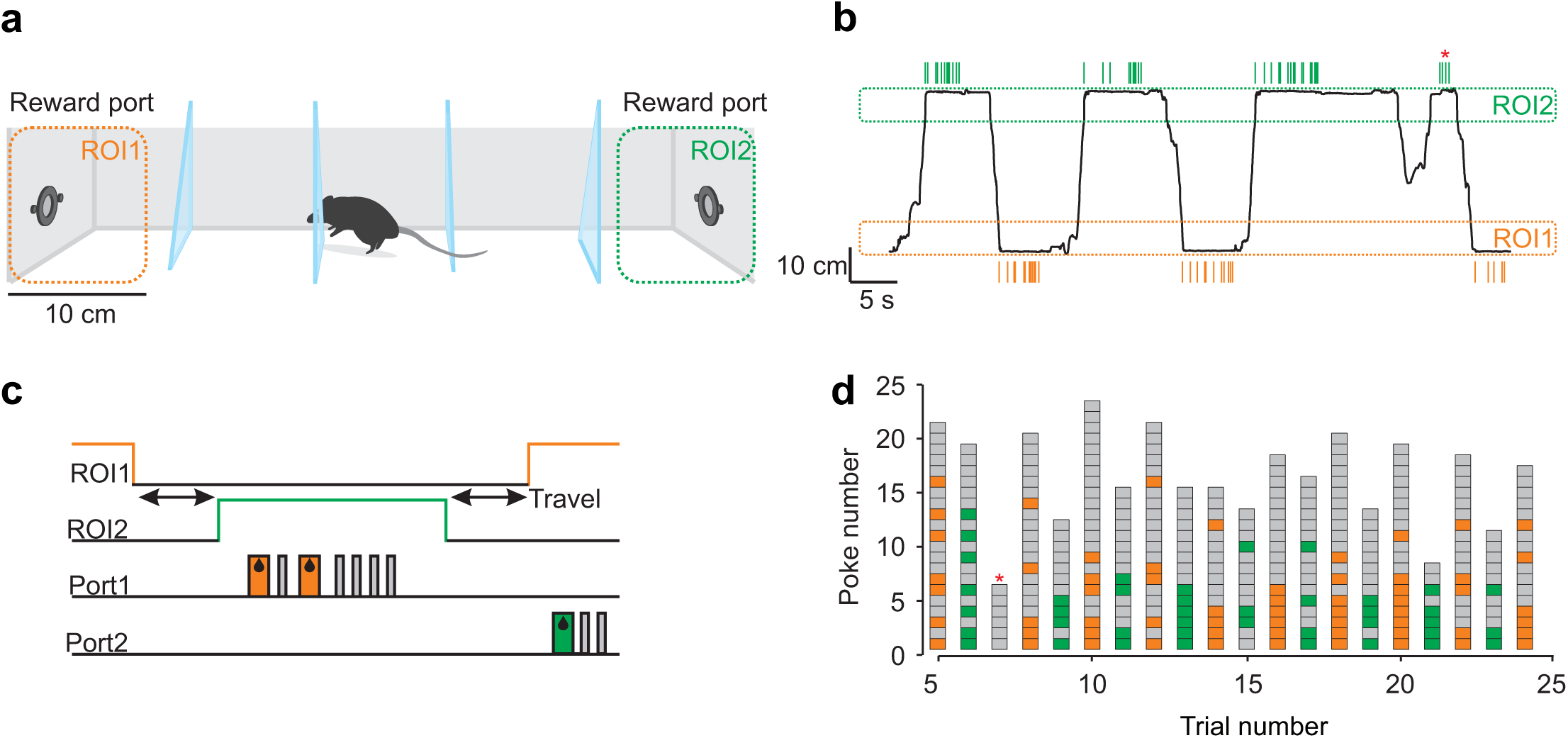
The probabilistic foraging task. (a) Schematic drawing of the foraging task apparatus. Mice shuttle back and forth between two reward sites, located at the opposite ends of an elongated box, to obtain water rewards. (b) Example snapshot of foraging behavior. The one-dimensional location of an example mouse along the long axis of the box is plotted as a function of time. The ROIs around each water port are marked as dashed rectangles, and green and orange ticks above and below the trajectory mark nose-pokes into the right and left ports, respectively. The red asterisk marks an error trial. (c) Task events during a single trial. Each trial starts with an exit from one of the ROIs. Following shuttling to the other end the mice would nose-poke multiple times and receive reward on some of the attempts on a probabilistic basis, before switching back. Green and orange rectangles mark rewards, grey rectangles mark omissions. (d) Example snapshot of foraging behavior, this time focusing only on the sequence of outcomes during nose-poking. Each column represents a single trial. Green/orange squares represent rewarded nose-pokes, gray squares – omissions. All trials in this example were correct except for the one marked with a red asterisk.

To incentivize goal-directed behavior, we introduced three trial types with different initial probability of reward (high-, medium- and low-quality patches), so that mice would benefit by taking into account the actual rewards received in a given trial, as opposed to adopting a fixed, reward-independent strategy. Reward probability decayed exponentially at the same rate regardless of initial probability (Fig. 2a; Eq. 1 in Methods). The three trial types were presented in a randomly interleaved manner, and were not cued to the mice. As expected, the average number of rewards gained was highest in high quality patches and lowest in low quality patches (Fig. 2b). However, the number of rewards mice obtained was lower than the maximum available. This is due to a trade-off between continuing to try to exploit the current port and leaving to try the other, at the cost of travel between ports (50 cm long, and 3.32 ± 1.70 s to cross from one side to the other (mean ± S.D.)). This tradeoff is formalized within optimal foraging theory by the marginal value theorem (MVT)^19^. This theorem states that optimal foragers should leave a depleting resource whenever the instantaneous reward rate within a patch drops below the average reward rate calculated across trials and taking into account travel times (Fig. 2c).

**Figure 2.**
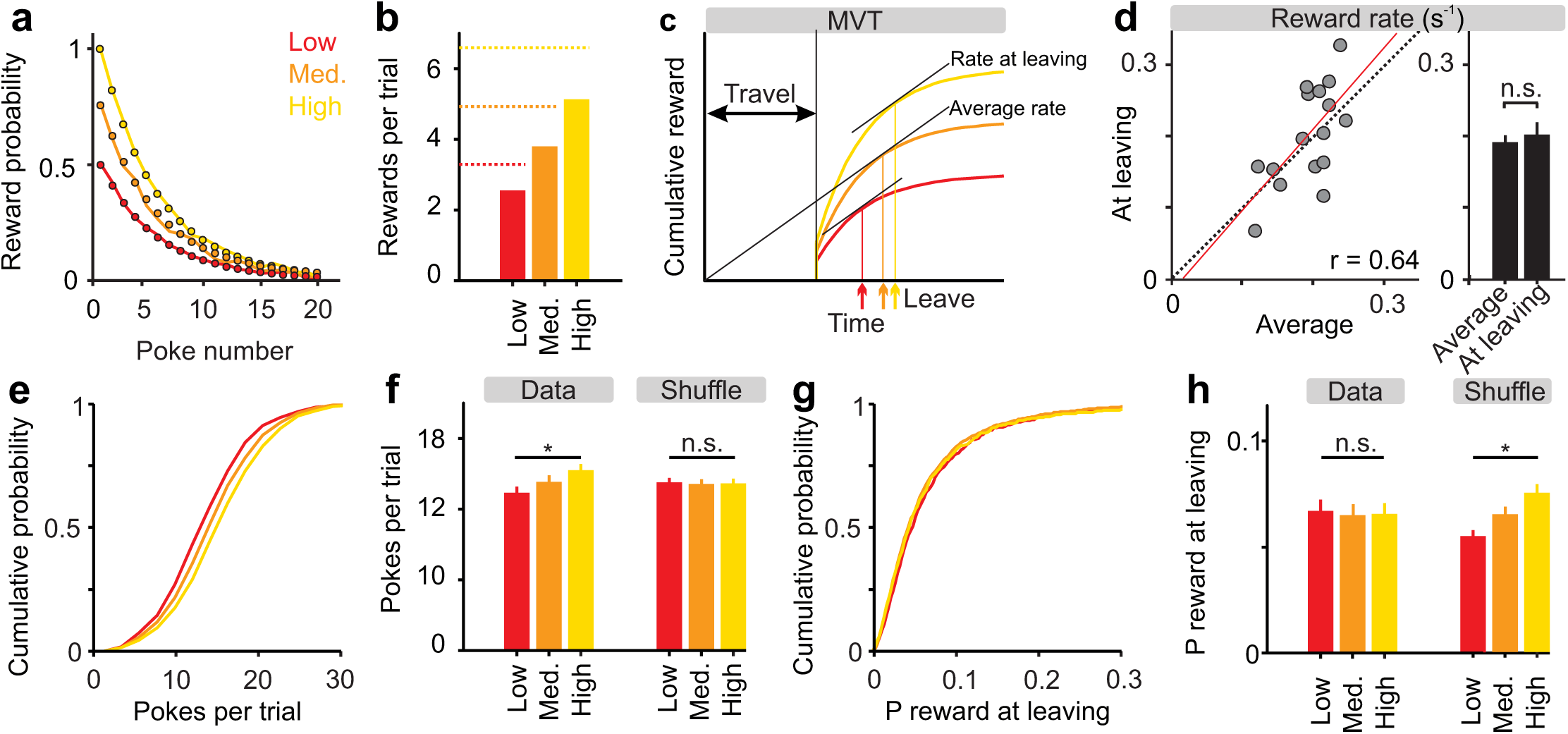
Reward statistics and task performance. (a) In each trial, reward probabilities were drawn from one of three exponentially decreasing functions, all sharing the same time constant but with a different scaling factor (1, 0.75 and 0.5), labeled high, medium and low, and color coded yellow, orange and red, respectively. Dots mark hypothetical values; solid lines are averages derived from data. (b) Bar plot showing the average number of rewards in each of the trial types (n = 16 mice). Dashed lines mark hypothetical maximal values, i.e the average number of rewards per trial, assuming the mice stay in place and poke indefinitely. (c) Schematic drawing of an optimal-agent’s behavior during foraging. According to the marginal value theorem, optimal leaving times are the times at which the instantaneous reward rate is equal to the average reward rate. When plotting the average cumulative reward (“medium” orange curve in this case) as a function of time from previous reward site exit, the average reward rate is the slope of the line that connects this curve at the time of leaving with the origin. Thus, the slope of this line is maximal when it is tangent to the curve. Consequently, better or worse trials result in later or earlier leaving times, respectively (vertical lines and arrows). (d) Left: Scatter plot of reward rate at leaving vs. average reward rate. Each circle represents one mouse (n = 16). Dashed line is the unity diagonal and red line is a linear regression curve, with its correlation coefficient shown as well (p < 0.001). Right: Bar plot showing the average reward rate at leaving and the average reward rate. There was no difference between the two, p > 0.05, Wilcoxon sign-rank test. (e) Cumulative distributions of the number of pokes per trial for the three trial types, averaged across mice (n = 16). (f) Bar plot showing the average number of pokes in each of the trial types. Bars on the left represent real data, and bars on the right represent shuffled data. Asterisks indicate significant difference between trial types (P < 0.05, ANOVA). (g) Cumulative distributions of the estimated reward probability after the last poke in a trial (i.e. at the time of switching) for the three trial types, averaged across mice (n = 16). (h) Bar plot showing the average estimated reward probability after the last poke. Bars on the left represent real data, and bars on the right represent shuffled data. Asterisks indicate significant difference between trial types, P < 0.05, ANOVA.

We tested the MVT in our data set and found that indeed the reward rate at the time of leaving was nearly identical to the average reward rate (0.19 ± 0.010 rewards·s^-1^ on average and 0.20 ± 0.017 rewards·s^-1^ at leaving (mean ± SEM); p = 0.54, Wilcoxon sign-rank test, n = 16 mice; Fig. 2D). Two additional predictions of the MVT are that^20^: (1) the mice should make more pokes in better than average trials, and less in worse than average trials; and (2) reward probability at the time of leaving should be the same regardless of the initial probability at the start of the trial. We tested these predictions in our data set by comparing the number of pokes made (Fig. 2e,f) and the expected reward probabilities at the time of leaving (Fig. 2g,h) for the three trial types. We found that indeed, number of pokes increased with increasing initial probability (F _(2,45)_ = 3.49, p = 0.04, one-way ANOVA, n = 16 mice), and that reward probabilities at the time of leaving were not significantly different across trial types (F _(2,45)_ = 0.049, p = 0.95, one-way ANOVA, n = 16 mice). In contrast, in a shuffled dataset, poke numbers were similar across trial types (F _(2,45)_ = 0.046, p = 0.95, one-way ANOVA, n = 16) and reward probabilities at the time of leaving varied with trial quality (F _(2,45)_ = 4.13, p = 0.022, one-way ANOVA, n = 16 mice; see Methods for details of the shuffling procedure). This confirms that the mice were sensitive to the statistics of individual port visits and suggests that the mice employed a near-optimal strategy in PFT.

### Optogenetic activation of DRN neurons during foraging prolongs active exploitation

We next examined the effect of activation of DRN 5-HT neurons in the PFT. Of the 16 mice examined, 10 expressed Cre-recombinase under the control of the SERT promoter (SERT-Cre) and 6 were wild-type litter-mates. Both groups of mice were infected in the DRN with a viral vector containing Cre dependent channelrhodopsin-2 (AAV2/9-Dio-ChR2-EYFP) and implanted with an optical fiber cannula above the site of infection (Fig. 3a)^21^. After 9 days of training, we started a 10-day testing period, in which we photostimulated randomly on 50% of the correct trials. Stimulation was triggered by the first nose-poke in each trial and lasted until the end of the trial (Fig. 3b). We found that the number of nose pokes in photostimulated trials was significantly greater than in control trials in ChR2-expressing mice (14.73 ± 0.55 vs. 13.61 ± 0.53 pokes per trial (mean ± SEM); p = 0.014, Wilcoxon signed-rank test, n = 10 mice). This difference was also significant comparing ChR2-expressing and wild-type mice (1.12 ± 0.36 vs. −0.46 ± 0.18 (mean ± SEM); p = 0.0047, Wilcoxon rank-sum test, n = 10 SERT-Cre and 6 wild-type mice; Fig. 3c-e). This effect was transient, occurring only on photostimulated trials but not on subsequent control trials (previous trial stimulated vs. previous trial non-stimulated: 13.67 ± 0.53 vs. 13.56 ± 0.53 pokes per trial (mean ± SEM); p = 0.56, Wilcoxon signed-rank test, n = 10 mice; Fig. 3c). Finally, the photostimulation effect was not dependent on patch quality (two-way ANOVA restricted to SERT-Cre mice, trial type: F_(2,54)_ = 4.28, p = 0.019, stimulation: F_(1,54)_ = 6.26, p = 0.016; interaction: F_(2,54)_ = 0.29, p = 0.75, n = 10 mice; Fig. 3f).

**Figure 3.**
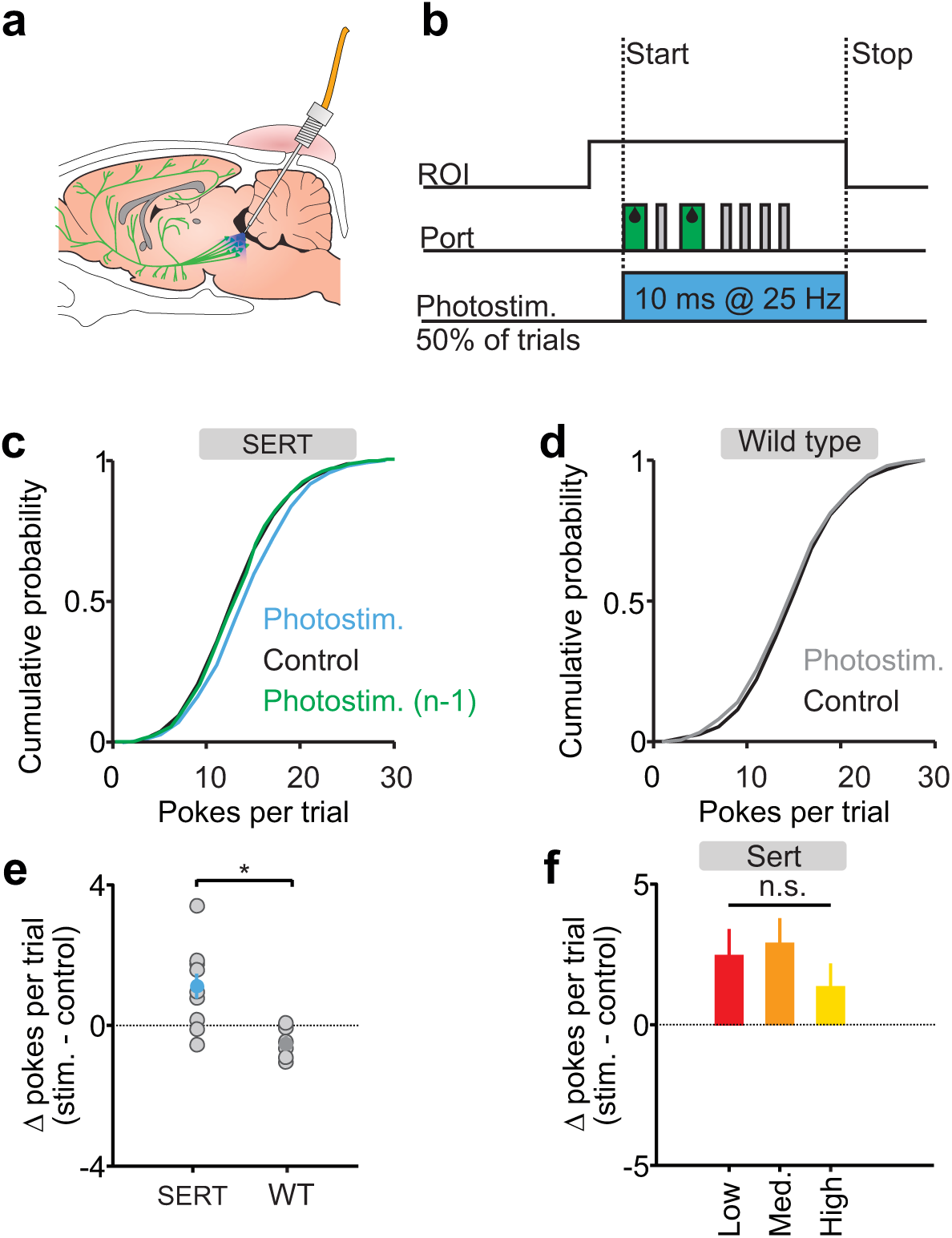
The effect of DRN 5-HT photostimulation on switching behavior. (a) Scheme of the locations of ChR2-YFP expression and optic fiber placement. (b) Schematic diagram of task events during a single trail, also showing the period of photostimualtion. Photostimulation was triggered by the first poke in 50% of correct trials and ended either when the mouse left the ROI or if 10 seconds had elapsed since the last poke. (c) Cumulative distributions of the number of pokes per trial for photostimulated trials (blue), control trials (black), and control trials immediately after photostimulation (green) averaged across the population of SERT-Cre mice (n = 10). (d) Cumulative distributions of the number of pokes per trial for photostimulated (blue) and control (black) trials, across the population of wild-type mice (n = 6). (e) Difference between average number of pokes in photostimulated and control trials for SERT-Cre (n = 10) and wild-type (n = 6) mice. Averages across mice are shown in filled circles. ∗p < 0.05, Wilcoxon rank-sum test. (f) Bar plot showing the average difference between average number of pokes in photostimulated and control trials for the three trial types in SERT-Cre mice (n = 10). There was no effect of trial type on this difference, P > 0.05, ANOVA.

### Optogenetic activation of DRN neurons biases leaving but not travel time

These results contradict the hypothesis that 5-HT promotes behavioral inhibition, as the willingness of mice to perform an active nose poking behavior was enhanced by optogenetic activation of DRN 5-HT neurons. We hypothesized that this phenomenon could reflect a bias in the process underlying the decision of whether to stay and poke again or leave after each nose poke. Consistent with this idea, we also found that photostimulation, which continued until the mouse exited the ROI, increased the delay to leave the foraging site, i.e. the interval between the last poke and ROI exit (control vs. stimulated trials: 3.84 ± 0.25 s in control vs. 4.26 ± 0.23 s in stimulated trials (mean ± SEM); p = 0.0039, Wilcoxon signed-rank test, n = 10 mice; Fig. 4b). However, once the mice had left the ROI, the time it took them to travel from one port to the other side was not affected (control vs. stimulated trials: 2.64 ± 0.23 vs. 2.77 ± 0.31 s (mean ± SEM); p = 0.56, Wilcoxon signed-rank test, n = 10 mice; Fig. 4c). However, the fact that photostimulation was not delivered during the travel period might explain the lack of effect on this measure. Therefore, we subsequently tested 6 of the 10 ChR2 expressing mice from our original cohort using a protocol in which a 2 second stimulation was triggered by ROI exit, so that stimulation took place during the travel period (Fig. 4a). In this case as well, no effect of photostimulation on travel time was observed (control vs. stimulated trials: 3.02 ± 0.31 vs. 3.19 ± 0.33 s (mean ± SEM); p = 0.22, Wilcoxon signed-rank test, n = 6 mice; Fig. 4d).

**Figure 4.**
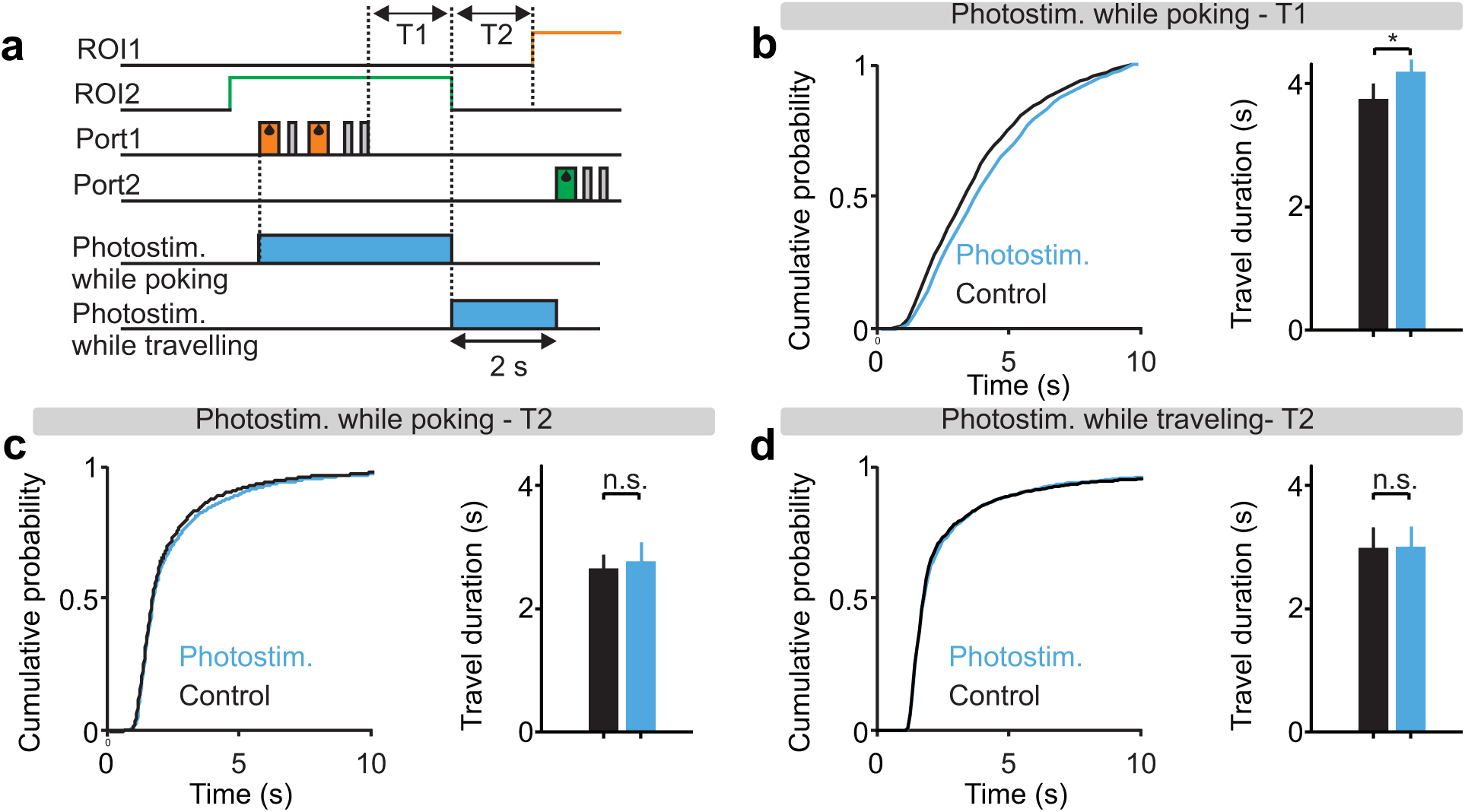
Lack of effect of DRN 5-HT photostimulation on travel duration. (a) Schematic diagram of task events during a single trial showing the period of photostimualtion. In the first protocol (photostimulation while poking) photostimulation was triggered by the first poke in 50% of correct trials, and in the second protocol (photostimulation while travelling) photostimulation was triggered by ROI exit in 50% of the correct trials and lasted for 2 seconds. We also defined two relevant time periods: (T1) time-to-leave - the interval between the last poke and ROI exit, and (T2) travel - the interval between ROI exit and subsequent ROI entry. (b) Left: Cumulative distributions of T1 durations for photostimulated (blue) and control (black) trials, averaged across the population of SERT-Cre mice (n = 10) in the photostimulation-during-poking protocol. Right: Bar plot showing the corresponding averages. ∗p < 0.05, Wilcoxon sign-rank test. (c) Left: Cumulative distributions of T2 durations for photostimulated (blue) and control (black) trials, averaged across the population of SERT-Cre mice (n = 10) in the photostimulation-during-poking protocol. Right: Bar plot showing the corresponding averages. p > 0.05, Wilcoxon sign-rank test. (d) Left: Cumulative distributions of T2 durations for photostimulated (blue) and control (black) trials, averaged across the population of SERT-Cre mice (n = 6) in the photostimulation-during-travelling protocol. Right: Bar plot showing the corresponding averages. p > 0.05, Wilcoxon sign-rank test.

### The proportional hazards model explains behavior in the probabilistic foraging task

Our analysis thus far suggests that DRN 5-HT stimulation biases the decision process that sets the tradeoff between continued nose-poking (‘exploitation’) and leaving the site (‘exploration’). We next sought to quantify this process more directly. Despite the agreement between our data and the predictions of the MVT (Fig. 2), we deemed it unlikely that the mice were using it directly for two main reasons: (1) the MVT requires an exact calculation of the expected reward probability after each nose-poke, which would require knowledge of the underlying task statistics, memory of the preceding sequence of rewards and omissions, and representation the three reward probabilities corresponding to the three trial types (Eq. 2 in Methods); (2) the MVT is deterministic in nature, and cannot account for the substantial variability we observed in mouse leaving times (this can be seen in the wide range of reward probabilities at the time of leaving in Fig. 2f; the MVT predicts these curves to be step functions)^22,23^. Indeed, several simpler heuristics have been proposed to explain decision-making in probabilistic foraging tasks^24^. Notably, a model known as the “proportional hazards” model^25^ not only aims at predicting mean leaving time, but also models a stochastic decision process that leads to leaving^20,26–28^. This model assumes that leaving decisions are taken randomly after each omission, based on an underlying hazard function (the conditional probability of leaving after n+1 misses, having experienced n consecutive omissions), and that the hazard function resets after each reward, albeit to a different value due to the influence of various predictor variables (Fig. 5a and Eq. 3, see Methods for further details of this model).

**Figure 5.**
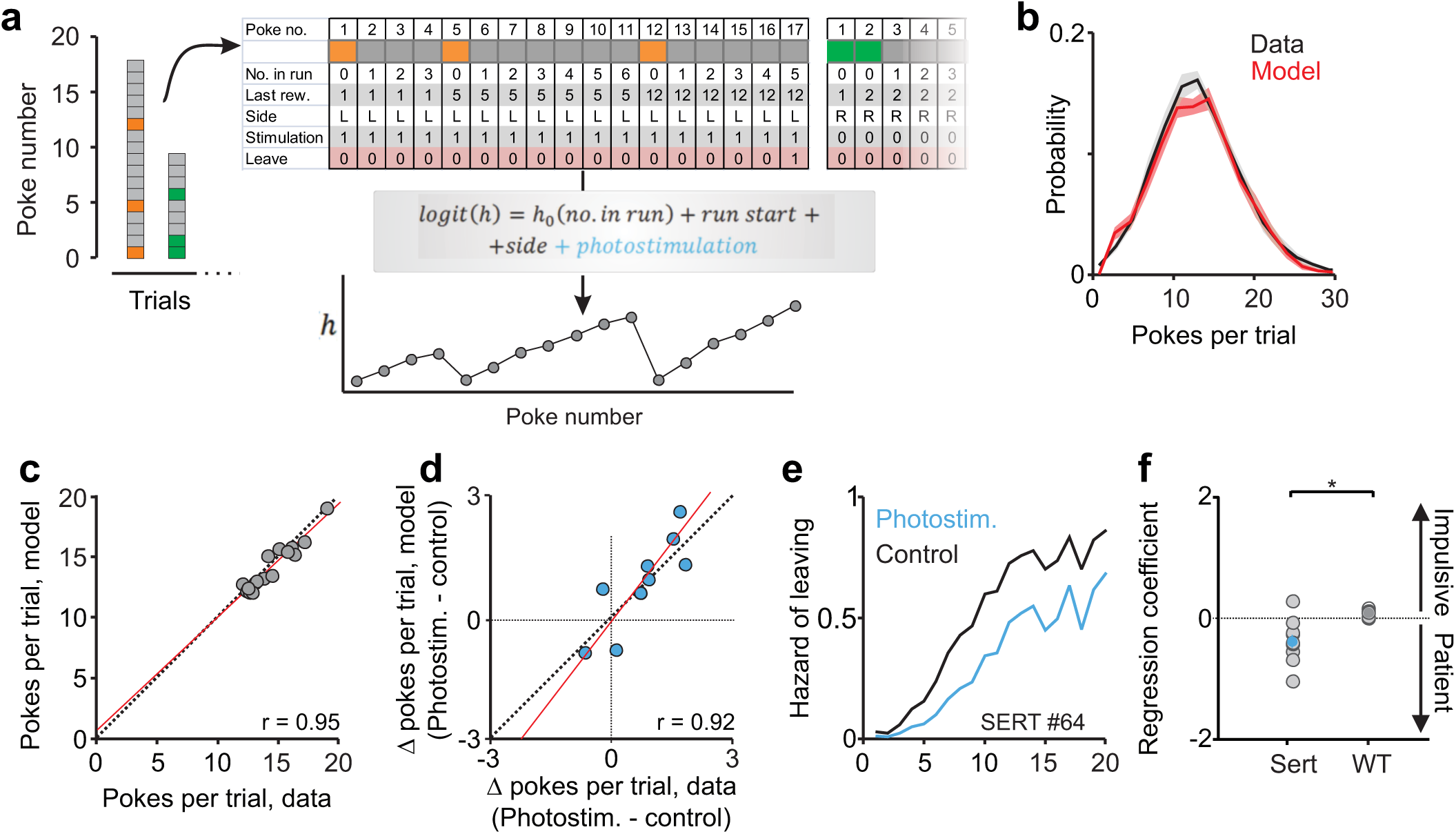
Fitting behavioral data using the proportional hazards model. (a) Schematic drawing of model-fitting pipe-line. Each nose-poke (pooled across trials and sessions for each mouse independently) was labeled with a vector of values corresponding to the different cox coefficients. These values, together with the outcome of each poke (stay or leave) were used to fit a logistic regression model - the outcome of which was an estimated hazard rate, h(n), that is reset at trial start and after each reward, and is multiplicatively changed by the different coefficient values. This hazard rate can be viewed as an estimate of the probability of leaving after each nose-poke and can therefore be used to simulate mouse leaving decisions. (b) Average pokes-per trial distributions for real and simulated data (n = 16 mice). (c) Scatter plot of simulated vs. real average number of nose-pokes per trial. Each circle represents one mouse (n = 16). Dashed line is the unity diagonal and red line is a linear regression curve, with its correlation coefficient shown as well (p < 0.001). (d) Same as (c) for the effect of photostimualtion on the number of nose-pokes per trial (p < 0.001). (e) The modelled hazard function for an example SERT-Cre mouse for phostostimulated and control nose-pokes. Note that decreased hazard means longer staying. (f) Cox regression photostimulation coefficient for SERT-Cre (n = 10) and wild-type (n = 6) mice. Averages across mice are shown in filled circles. ∗p < 0.05, Wilcoxon rank-sum test.

To test how well the proportional hazards model captures individual mouse behavior, we used half the data to fit model parameters and cross-validated it using the second half. These simulations were in excellent agreement with our data, capturing both the mean and variability of the leaving time distributions (Fig. 5b). Furthermore, the model was accurate enough to predict idiosyncratic individual differences in poke numbers across mice (Fig. 5c), as well as the effect of photostimulation on a mouse-by-mouse basis (Fig. 5d). This model allowed us to examine the effect of photostimulation on the estimated hazard rate, expressed as the Cox regression coefficient for that variable. We found it to be significantly negative (that is, divisive or patience promoting) in SERT-Cre mice (p = 0.014, Wilcoxon signed-rank test, n = 10 mice), and significantly lower than that of wild-type mice (p = 0.0075, Wilcoxon rank-sum test, n = 10 SERT-Cre and 6 wild-type mice; Fig. 5e,f).

### Vigor of nose-poking continuously reports the hazard rate of leaving

The estimated hazard function in the proportional hazards model can be interpreted as a latent decision variable – the instantaneous propensity to switch foraging sites. This variable plays a similar role to that of accumulated evidence in integration-to-bound decision models^29^. In this interpretation, the effect of optogenetic activation of DRN 5-HT neurons is to slow down the dynamics of the decision variable. A shortcoming of this interpretation is that it is an indirect one – it infers the dynamics of the decision variable only by extrapolating backwards in time from the decision process’s termination point. We therefore wondered whether a more direct readout during the unfolding of the decision process might be available. To address this question, we examined the correlation between different behavioral variables related to nose-poking and the latent decision variable in the proportional hazards model (i.e. the estimated hazard of leaving; Supplementary Fig. 2). We found that the duration of nose-pokes for omitted rewards and the duration of inter-poke-intervals appeared to mirror the value of this hypothetical decision variable (Supplementary Fig. 2; note that according to our model, rewards reset the decision variable, which may explain the lack of correlation between the duration of rewarded nose-pokes and the hazard of leaving). Using these two variables, we calculated a measure that we refer to as the “omission duty cycle” (ODC; Fig. 6a). The ODC, which essentially measures the rate or vigor of nose poking, was indeed strongly **negatively** correlated with the model hazard rate (Fig. 6b).

**Figure 6.**
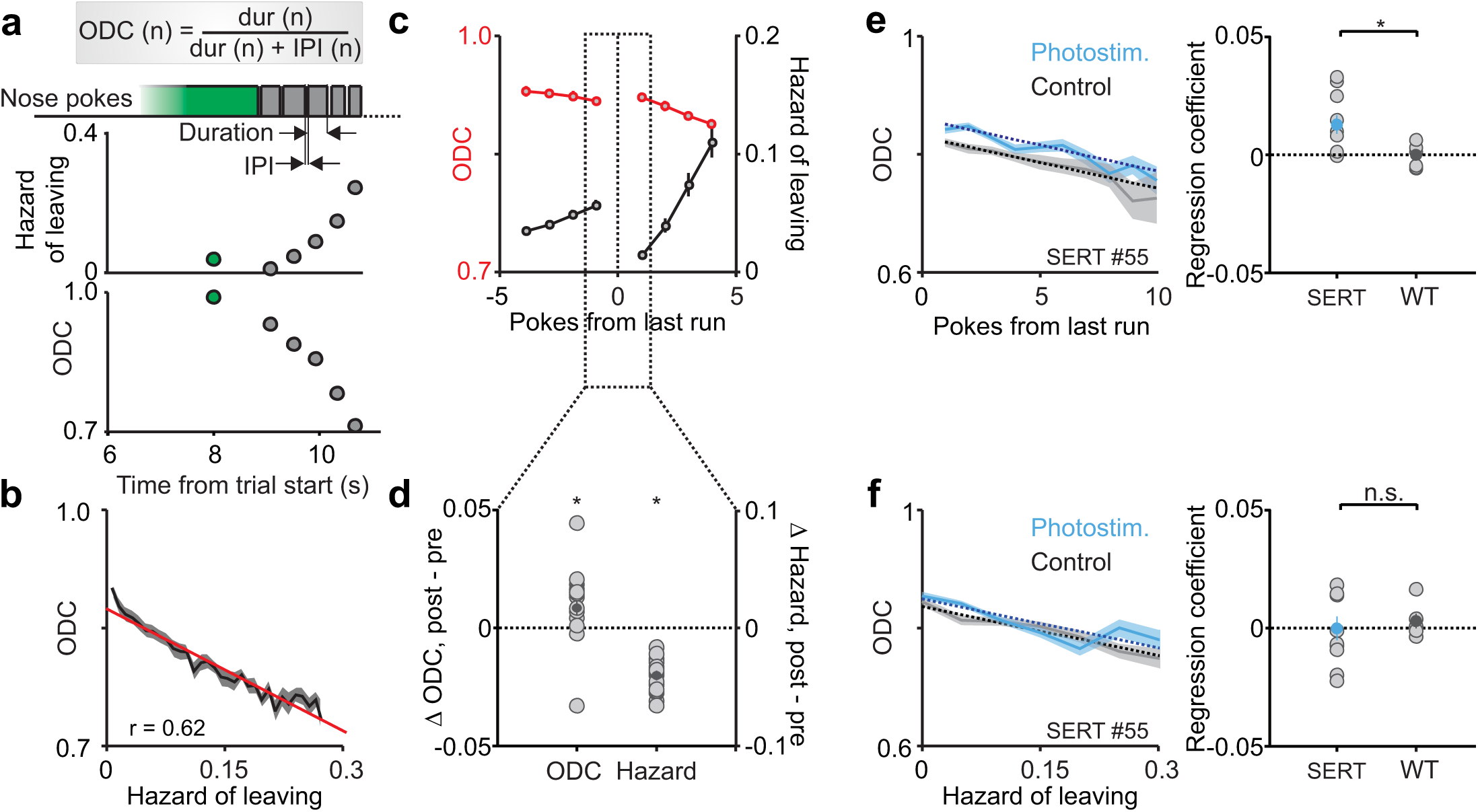
The effect of DRN 5-HT photostimulation on the microstructure of behavior. (a) Example trial represented in real time and starting from the last reward in that trial. Top: Each rectangle represents a single nose-poke (green marks the last reward and grey, the subsequent omissions). Middle: estimated hazard for each nose-poke. Bottom: omission duty cycle (ODC). (b) Correlation between ODC and estimated hazard. The red line is a linear regression curve, with its correlation coefficient shown as well (p < 0.001). (c) Rewards reverse an overall (across trial) monotonic decrease in ODC (red) and an overall increase in hazard (green). The plot shows the average ODC and hazard as a function of poke number, aligned on the last reward and averaged across the population of mice (n = 16). Note that despite a decreasing (increasing) trend, the ODC (hazard) immediately after the last reward is higher (lower) than the one just before it. ∗p < 0.05, Wilcoxon sign-rank test. (d) Difference between average ODC and hazard immediately before the last reward compared to immediately after it across the population of mice (n = 16). ∗p < 0.05, Wilcoxon sign-rank test. (e) Left: The ODC for an example SERT-Cre mouse for phostostimulated and control nose-pokes aligned on last reward. The dashed lines are a linear regression curves, (p < 0.001 for the phostostimulation coefficient). Right: Regression photostimulation coefficients for SERT-Cre (n = 10) and wild-type (n = 6) mice. Averages across mice are shown in filled circles. ∗p < 0.05, Wilcoxon rank-sum test. (f) Left: The ODC for the same SERT-Cre mouse shown in (D) for phostostimulated and control nose-pokes as a function of estimated hazard. The dashed lines are a linear regression curves, (p > 0.05 for the phostostimulation coefficient). Right: Regression photostimulation coefficients for SERT-Cre (n = 10) and wild-type (n = 6) mice. Averages across mice are shown in filled circles. p > 0.05, Wilcoxon rank-sum test. (g) Left: Inter-poke-interval (IPI) as a function of nose-poke number within a trial. Right: correlation between IPI and estimated hazard. The red line is a linear regression curve, with its correlation coefficient shown as well (p < 0.001).

To further relate the ODC measure to the decision-making process, we noted that while the overall pattern is of monotonic increase in the hazard of leaving and a monotonic decrease in ODC throughout trials, this pattern is reversed when aligning on the last reward due to the resetting effect of rewards, such that the hazard is higher immediately before the last reward compared to immediately after it (0.056 ± 0.0041 before and 0.014 ± 0.0023 after the last reward (mean ± SEM); p = 0.00044, Wilcoxon signed-rank test, n = 16 mice; Fig. 6c,d). We found that, as predicted, this reversal was true for the ODC as well (0.86 ± 0.010 before and 0.87 ± 0.011 after the last reward (mean ± SEM); p = 0.0084, Wilcoxon signed-rank test, n = 16 mice; Fig. 6c,d). These results provide strong evidence that the vigor with which mice nose-poke is: (1) monotonically decreasing in a manner reflecting the decreasing reward probability; and (2) non-trivially correlated with the effect of rewards on the modelled hazard function. The latter observation argues against the possibility that simpler variables that would be expected to evolve monotonically during nose-poking, such as muscle fatigue, could account for the observed relationships.

We next examined the effect of DRN 5-HT photostimulation on ODC. Since stimulation decreases the hazard of leaving (Fig. 5) and ODC is negatively correlated with the hazard rate, we predicted that the ODC would be higher in photostimulated vs. control trials. Using linear regression analysis, we found that photostimulation indeed increased the ODC of SERT-Cre mice (p = 0.0098, Wilcoxon signed-rank test, n = 10), and that this increase was significantly higher in SERT-Cre compared to wild-type mice (p = 0.011, Wilcoxon rank-sum test, n = 10 SERT-Cre and 6 wild-type mice; Fig. 6e). Furthermore, we reasoned that if the ODC truly reflects the modelled hazard function, then it ought to be the same in photostimulated vs. control trials, when conditioned on the hazard function (since this quantity already takes into account the effect of photostimulation on the decision process). Using similar linear regression analysis, we found that, when plotting the ODC as a function of the modelled hazard, photostimulation had no effect on the ODC of SERT-Cre mice (p = 0.90, Wilcoxon signed-rank test, n = 10), nor was there a difference between SERT-Cre compared to wild-type mice (p = 0.54, Wilcoxon rank-sum test; Fig. 6f).

These results show that the vigor of poking behavior is closely linked to the underlying decision-making process and can be used as a real-time read-out of a latent decision variable^30^. Furthermore, the effects of 5-HT on foraging behavior were well explained by a model in which it biases decisions by multiplicatively scaling the latent decision variable on which leaving decisions are based.

## Discussion

In this paper, we found that optogenetic activation of DRN 5-HT neurons during foraging promotes active exploitation of the current foraging site, increasing the time to switch. This result in some ways resembles the previous finding that DRN 5-HT activation increases the ability to wait for delayed outcomes^15,16^. However, because in this task “waiting” required active behavior (nose poking) the results argue strongly against the hypothesis that 5-HT enhances waiting by enhancing behavioral inhibition. Rather, 5-HT appears to stabilize an on-going behavior by reducing the probability (or rate) at which animals switch to the next one - here, stabilizing nose-poking while reducing the probability of leaving. This behavior could be well described by a simple model, the proportional hazards model^20,26–28^. In this model, the effect of 5-HT stimulation was to proportionately scale down the hazard rate, thereby reducing the probability to leave. Thus, 5-HT appears to act as multiplicative bias on the latent decision variable in the foraging decision process.

### Serotonin biases decision-making by scaling a latent decision variable

Foraging behavior, namely the searching for an accumulation of food and other resources, is fundamental to any animal’s survival. Natural environments impose significant constraints on this behavior, forcing animals to balance various consideration in order to achieve optimal results^31^. Of particular interest for the theory of optimal foraging is the exploration-exploitation dilemma: should the animal stay and exploit a depleting resource, or leave instead to forage elsewhere? Within this framework, the MVT describes the optimal strategy, which is deciding to leave at the point in time when the instantaneous rate of rewards drops below its average value^19^. However, the MVT, as originally formulated, applies to deterministic environments, where foragers have perfect knowledge of the instantaneous rate and a complete memory of the average rate. Although these assumptions are not realistic, and the solution for the stochastic case is quite intricate^22^, a key prediction of optimality remains relevant: switching decisions should be made when instantaneous reward probability is equal to some fixed value, irrespective of whatever has happened up to this point^20,27^. Our study confirms that mice follow this prediction in a laboratory version of a foraging scenario.

In order to model the dynamics of the stochastic environment, we used a simple heuristic decision rule, the proportional hazards model^25^. We found that this model captures not only the fixed relationship between instantaneous and average reward rate at leaving time, but also the variability of this decision point across the mouse population (Fig. 5c) and the case-by-case effects of photostimulation across mice (Fig. 5d). Here, the effect of DRN 5-HT activation by photostimulation was to decrease the hazard rate in a proportional manner. It thereby increased the average time to reach decision threshold, increasing the time spent nose-poking before deciding to leave.

The proportional hazards model shares commonalities with models that have been widely used in related fields, such as interval timing^32^ and perceptual decision-making^29^. In this class of models, before reaching a decision, agents integrate information in the form of elapsed time^33,34^ or noisy samples of sensory input^29,35^. The amount of accumulated information during the accumulation process is a latent or hidden variable that, through behavioral studies, can only be indirectly inferred through modeling the timing of overt behavior: the timing and type of choices made. An important source of evidence in favor of such models has been provided by the ability to find correlates of decision variables in neural activity patterns recorded in brain regions hypothesized to underlie the decision process^29,35,36^. Here, we found that a dynamic and readily observable microfeature of mouse behavior, the vigor of nose-poking during foraging, strongly correlated with the latent decision variable in the proportional hazards model. This observation offers significant support for the validity of the proportional hazards model for foraging. It also supports the notion that readily observable dynamic microfeatures of behavior reflect latent decision variables, providing general insight into the “covert” processes underling “overt” decisions^37,38^.

Our success in modeling the effect of 5-HT using a simple decision model encourages us that a computationally-based and behaviorally-constrained approach will be a valuable way to approach the interpretation of 5-HT function. At the same time, our results leave open the question of how more precisely to interpret the influence of 5-HT, since the decision-variable that was modulated is highly abstract and represents the conjunction of many more specific factors that might enter into the decision to stay or leave. Broadly speaking, these fall into two categories. First, there are factors that represent the calculation of the costs and benefits of decision options. These include the cost or effort (‘handling cost’ in the foraging literature) of foraging at a given site (including the perceived speed at which time elapses), the expected reward gained for each foraging attempt and the expected cost of travel to the next foraging site. Second, there are factors related to the uncertainties inherent in the probabilistic foraging task, especially the uncertainty related to the stochastic nature of reward delivery^31^.

For an optimal Bayesian agent, estimating the costs and benefits of decision options must take into account the uncertainties of those estimates. Thus, the impact of 5-HT on the hazard rate of leaving might be due to a more direct effect on costs or benefits of staying vs. leaving, or it might arise from a direct modulation of uncertainty which only indirectly impacts estimates of cost and benefit^39^. We currently favor the latter interpretation for two reasons. First, a growing set of evidence suggests that 5-HT does not exert the sort of motivational biases that arise from natural rewards or punishments^16,40^. If 5-HT modulated the costs or benefits of on-going events with which it was correlated, one might expect this to be revealed in a broad range of situations. Yet, 5-HT does not produce place preference or bias thigmotaxis in the open field^41^ and 5-HT stimulation applied during the outcome of a two-alternative choice value-based decision-task, does not bias choices^16^. Second, the endogenous activity of 5-HT neurons reports unexpected events regardless of valence, as appropriate for a signal related to uncertainty^42^.

One limitation of the present study is that the task was too complex to be modeled precisely by a simple model; the proportional hazards model is not the optimal solution to this task and the complexity of the optimal solution suggests that it is not likely used by mice. This issue could be addressed in future studies using refined tasks in which simple optimal solutions exist. This, together with additional manipulations of task variables, should help to ascertain more precise interpretation of the influence of 5-HT.

### Relationship to serotonin effects in other tasks

We have interpreted our results as arguing against one of the more prominent theories of 5-HT action: the behavioral inhibition hypothesis^3^. When it was first suggested that 5-HT causes behavioral inhibition, it was also noted that its effects are context-dependent. Namely, 5-HT is important primarily in those situations where a choice needs to be made between active response and passivity, and it biases choices towards the latter. This idea has been developed in a variety of studies exploring the linkage between 5-HT and patient waiting for delayed rewards. Recordings showed that DRN neurons (possibly, but not conclusively 5-HT neurons) increase their firing rates during waiting, and that trial-by-trial fluctuations in their firing rates were predictive of impulsive leaving^14^. Moreover, optogenetic activation of DRN 5-HT neurons prolongs waiting^15,16^. Similarly, fiber photometric imaging of DRN 5-HT neurons during a foraging-like task has shown increased activity specifically during nose-poking and reward consumption and lower activity during travel^43^.

Here, we have shown that optogenetic activation of DRN 5-HT neurons prolongs the willingness of mice to forage for stochastically-delivered rewards. There are apparent similarities and differences between this foraging task and waiting tasks that may help to provide some deeper insight into the function of 5-HT in this context. First, in terms of similarities, we note that waiting can be considered a sort of foraging situation in which rewards might be expected but are not forthcoming. Indeed, in the foraging task, the period after the last reward poke, during which a mouse continues to attempt to gain rewards for some time, is almost precisely equivalent to a waiting task, albeit an active one (because nose pokes are required). Furthermore, the same class of integrate-to-threshold models can be used to model both foraging and waiting tasks^32,33,34^. Thus, it is not entirely surprisingly that 5-HT effects on waiting and foraging are both to prolong the duration.

On the other hand, the key difference between the probabilistic foraging task and previously-used waiting tasks is that in waiting tasks, passivity and waiting are coincident: the active choice is to leave and the passive choice is to stay and wait. In contrast, in the foraging task, waiting and passivity are decoupled: the choice is no longer between passive and active behaviors, but between two different types of active behavior, nose poking or locomoting. Thus, we could distinguish between two previously confounded interpretations. The reason that 5-HT favors patient waiting is not because it favors behavioral inhibition or passivity but because it favors persistence in a current behavior, even if it is active. One possible interpretation of this finding is that 5-HT does simply that: it favors persistence by making behavioral transitions (switching from one behavior to another) less likely. This interpretation can explain the observation that DRN photostimulation does not bias travel times between reward sites when stimulation occurs after the mouse is already in transit (Fig. 4 and ref. 41). It would also be consistent with an observed increase in active escape behavior (reduction of immobility) in the forced swim test that is induced by stimulation of medial prefrontal cortex axons in the DRN^44^ (although it is not known whether this stimulation activated or inhibited DRN 5-HT neurons^45^). However, as discussed above, another interpretation is that these effects are instead the consequences of effects of 5-HT on one or more factors that contribute to the decision variable on which the behavioral transitions are based. For example, by increasing the perceived uncertainty of the instantaneous estimates of the reward rate, 5-HT could bias decisions in favor of continued nose-poking in the PFT^39^.

Despite the relative similarity and even possible consistency of the above results, one should not lose site of the complexity of the 5-HT literature, which implicates this molecule in a host of functions with little obvious relationship to either foraging, waiting or persistence. However, the application of simple and relatively abstract computational models, such as the one used here, may provide a useful way toward a more satisfactory overarching understanding. Rather than seeing 5-HT’s effects in terms of a suite of diverse behaviors for which the overarching behavioral theme is missing, one may look to commonality of action at a lower mechanistic level. Thus, a class of model functions, such as integration-to-bound, may be considered generic algorithms that are reused in a diverse array of computations to implement widely divergent behaviors. If the action of 5-HT is best described at this algorithmic level, say as a way of modulating integration rate in the circuits that implement integration-to-bound, then the range of behaviors in which 5-HT acts will be as diverse as the instances in which this algorithm is applied. For example, in larval zebra fish 5-HT is involved in modulation of short-term locomotor memory to change visuo-motor gain^46^. While the relationship between sensorimotor gain adaptation and probabilistic foraging is obscure at the behavioral level, neural integration is a common component of both. The investigation of 5-HT action at the level of postsynaptic target circuits will be important to test such hypotheses.

## Materials and methods

### Animal subjects

Mouse lines, surgical procedures for virus injections and optic fiber cannula implantation, and optical setups for optogenetic stimulation were identical to those described in previous papers from our lab^21^.

Sixteen adult male C57BL/6 mice (10 SERT-Cre mice and 6 wild-type littermates) were used in this study. All experimental procedures were performed in accordance with the Champalimaud Centre for the Unknown Ethics Committee guidelines, and approved by the Portuguese Veterinary General Board (*Direcção-Geral de Veterinária*, approval 0421/000/000/2016). The SERT-Cre mouse line^47^ was obtained from the Mutant Mouse Regional Resource Centers (stock number: 017260-UCD). The mice were kept under a normal 12 hour light/dark cycle, and training as well as testing occurred during the light period. During training and testing the mice were water deprived, and water was available to them only during task performance. Food was freely accessible to the mice in their home cages.

Behavioral training started 2-10 weeks after virus injection and lasted for 9 days, at the end of which we commenced testing (experimentation). Testing periods consisted of 10 consecutive daily sessions. 10 of the 16 mice were only tested once, and 6 SERT-Cre mice (out of the 10) were tested again using a different photostimulation protocol (see below). This second testing period started one month after the end of the first one. The experimenters were blind as to the mice’s genotype throughout training and testing periods.

### Stereotaxic adeno-associated virus injection and cannula implantation

Mice were anesthetized with isoflurane (4% induction and 0.5 – 1% for maintenance) and placed in a stereotaxic frame (David Kopf Instruments, Tujunga, CA). Lidocaine (2%) was injected subcutaneously before incising the scalp. In order to infect serotonergic neurons with Channelrhodopsin-2 (ChR2) a craniotomy was drilled over the cerebellum and a pipette filled with a viral solution (AAV2.9.EF1a.DIO.hChR2(H134R)-eYFP.WPRE.hGH, 10^13^ GC/mL, University of Pennsylvania) was lowered to the DRN (Bregma −4.7 AP, −2.9 DV) with a 32° angle toward the back of the animal. The viral solution (1 µL) was injected using a Picospritzer II (Parker). After waiting for 10-15 minutes the pipette was removed from the brain and an optical fiber (200 µm core diameter, 0.48 NA, 4-5 mm long, Doric lenses) was lowered through the same craniotomy such that its tip was positioned 200 µm above the injection point. The implant was cemented to the skull using dental acrylic (Pi-Ku-Plast HP 36, Bredent, Senden, Germany). Mice were monitored until recovery from the surgery and returned to their home cages. Gentamicin (48760, Sigma-Aldrich, St. Louis, MO) was topically applied around the implant.

### Optogenetic stimulation

In order to optically stimulate ChR2 expressing 5-HT neurons we used blue light from a 473 nm laser (LRS-0473-PFF-00800-03, Laserglow Technologies, Toronto, Canada or DHOM-M-473-200, UltraLasers, Inc., Newmarket, Canada) that was controlled by an acousto-optical modulator (AOM; MTS110-A1-VIS or MTS110-A3-VIS, AA optoelectronic, Orsay, France). Light exiting the AOM was focused into an optical fiber patchcord (200 µm, 0.22 NA, Doric lenses), connected to a second fiber patchcord through a rotary joint (FRJ 1x1, Doric lenses), which was then connected to the chronically implanted optic fiber cannula.

### Histology

In order to confirm successful viral expression of ChR2-eYFP and optical fiber placements we used post-mortem histology at the end the experiments. Mice were deeply anesthetized with pentobarbital (Eutasil, CEVA Sante Animale, Libourne, France) and perfused transcardially with 4% paraformaldehyde (P6148, Sigma-Aldrich). The brain was removed from the skull, stored in 4% paraformaldehyde overnight and kept in cryoprotectant solution (PBS in 30% sucrose) for one week. Sagittal sections (50 mm) were cut in a cryostat (CM3050S, Leica, Germany), mounted on glass slides with mowiol mounting medium (81381, Sigma-Aldrich, St. Louis, MO). Scanning images for YFP and transmitted light were acquired with an upright fluorescence microscope (Axio Imager M2, Zeiss, Oberkochen, Germany) equipped with a digital CCD camera (AxioCam MRm, Zeiss) with a 5X or 10X objective. In a previous study using the same Cre-dependent optogenetic approach and the same mouse line we reported that 94% of ChR2-eYFP positive neurons were serotonergic^21^.

### The probabilistic foraging task

16 water-deprived mice were trained in a probabilistic foraging task. The apparatus was an elongated 50X14 cm chamber with two water-reward ports, one at each end. Each port had an infrared emitter/sensor pair located on its sides to measure nose-pokes (model 007120.0002, Island motion corporation) and a metallic tube running through its center for water delivery (controlled by a valve - LHDA1233115A, The Lee Company, Westbrook, CT). Four partitions were placed at regular intervals along the chamber. Each partition blocked about one half of the corridor’s width, forcing the mice to zig-zag between them when crossing from side to side. All task-related events were controlled using a behavioral control system, Bcontrol, that was developed by Carlos Brody (Princeton University) in collaboration with Calin Culianu, Tony Zador (Cold Spring Harbor Laboratory) and Z.F.M.

Each trial consisted of a sequence of nose-pokes in one of the two reward ports. Each nose-poke was rewarded with some probability by a 3ul drop of water. Reward probabilities decreased after each nose-poke, forcing the mice to alternate between sides during the session. Correct trials were ones in which the mice alternated sides, and error trials were ones in which the mice returned to the same reward port without nose-poking in the other one. Reward probabilities in each correct trial were drawn from one of three, equally likely exponentials, each decreasing with poke number

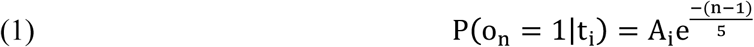

Where t_i_ is the i^th^ trial type (i = 1,2,3) corresponding to low, medium and high quality trials. These types differed in their exponential scaling factors, such that A_1_ = 0.5, A_2_ = 0.75, A_3_ = 1. N marks the poke number within a trial, and o_n_ is the outcome of the n^th^ poke (1 for reward and 0 for omission). Trial types (exponential scaling factors) were randomly interleaved and trial type identity was not cued to the mice. Reward probability was set to zero during error trials. For technical reasons, the reward probability was set to zero after the 20th nose-poke, this led to only a marginal deviation from true exponentials.

The mice were tracked on-line using Bonsai, a visual programming framework^48^, and their position was used to control trial transitions. We defined a 10 cm square region of interest (ROI) around each port. ROI exits signaled the end of one trial and the beginning of the next one. Additionally, to avoid exceptionally long photostimualtion durations, trials were terminated if 10 seconds had elapsed since the last nose-poke exit, even if the mouse was still in the ROI.

Each testing session lasted about 30 minutes in which the mice performed at least 80 trials, gaining an average of 3.75 rewards per trial. We found that the behavior of the mice was very consistent throughout the session. However, some trials, particularly towards the end of the session, were unusually short, thus making the distribution of pokes numbers per trial bimodal (Supplementary Fig. 1). We therefore consider for analysis only the first 60 trials in each session, and only those trials that contained more than two pokes in them.

### Photostimulation protocols

In this paper, we present the results of two experiments. The behavioral task was identical in both (as described above), yet the photostimualtion protocol differed between the two. In the first experiment, photostimulation (a train of 10 ms wide pulses at 25Hz, measuring 5mW at fiber tip) was triggered by the first nose-poke in a trial and lasted until the trial’s end (i.e. by either ROI exit, or if 10 seconds had elapsed since the last nose-poke exit). In the second experiment a similar photostimulation train was triggered by ROI exit and lasted for 2 seconds. 16 Blue LEDs were placed inside the box, 8 above each port, and delivered a flickering masking light, to prevent behavioral changes due to the mere laser light flashes during photostimulated trials. The masking light was identical to the photostimulation in frequency and duration and was present in all trials.

## Data analysis

All data analysis was performed using custom-written software in MATLAB (Mathworks, Natick, MA). In all figures, average data and error bars, or shaded patches around curves, represent mean ± S.E.M.

### Subjective reward probability

In the analysis shown in Fig. 2 we calculated the reward probability at the time of leaving (i.e the expected reward probability in the next poke, if it were to be made). While the actual probabilities are given by Eq. 1, trial types were not cued to the mice, making it impossible to know exactly each poke’s reward probability. Instead, we considered the most accurate estimate, assuming perfect knowledge of the task’s structure and reward history leading up to a nose-poke. The probability of gaining a reward in poke n+1 given trial history o_1_,…, o_n_ is thus:

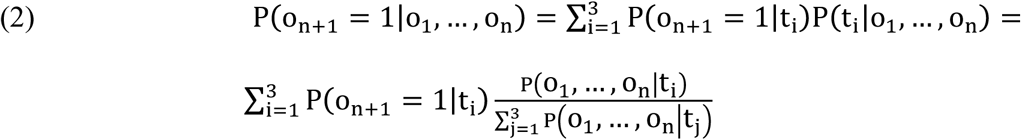

Where t_i_ is the i^th^ trial type (i = 1,2,3) and 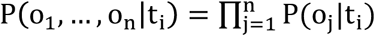is calculated using Eq. 1. In Fig. 2d we use this quantity to obtain the instantaneous reward rate by calculating, for each nose-poke, the ratio between this and the poke’s duration.

### Shuffled behavioral data

In Fig. 2 we also compare real to shuffled datasets. Shuffling was done individually for each mouse by first considering all potential trials, namely the 20 poke-long sequences of rewards and misses that are calculated using equation 1. Next, we paired each such potential trial with a randomly chosen trial length, selected from the actual data. This resulted in a shuffled data set in which average trial lengths were preserved while all other correlations between trial reward structure (such as trial type) and behavior were destroyed.

### Travel time analysis

In Fig. 4 we analyze the effect of photostimulation on travel times. Since the travel time is not well-defined in error trials (when the mice leave and return to the same location without visiting the other reward port in-between), we considered for this analysis only those trials in which both current and subsequent trials were correct. Additionally, since photostimulation terminated if 10 seconds had elapsed since the last nose-poke exit, even if the mouse was still in the ROI, such trials were excluded from this analysis as well. Consistent with the result shown in Fig. 4b, the fraction of such trials was higher in the photostimualtion condition (data not shown).

### Cox proportional hazards regression model

In order to model individual mouse choice behavior and its modulation by various factors (e.g. DRN photostimualtion), we used the Cox proportional hazards regression model. This semi-parametric model calculates the probability of leaving after each poke according to a baseline hazard (the probability of leaving immediately after a poke as a function of its number) which is estimated from the data, and that may be changed multiplicatively by trial-general covariates. The model is reset after each reward, but potentially to a different value, depending on the covariates. Therefore, we fitted leaving probabilities for all nose-pokes as a function of their distance from the previous reward rather that from trial start. To do so, each trial was segmented into one or more sub-trials, or runs. Each such run was bracketed from the left by a reward (except for the first one, in cases were the first poke in a trial was an omission) and from the right (except for the last one, in the very common cases were the last poke in a trial was an omission). Note that length zero runs were also allowed, if two rewards were delivered consecutively. As noted in ref. 25, in the discrete case this model is equivalent to a logistic model of the form:

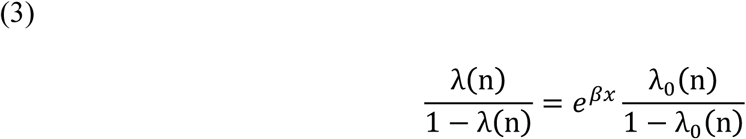

Where λ(n) represents a hazard function (hazard rate of leaving after the n^th^ poke. λ_0_(n) represents a baseline hazard function, that is the hazard function when all the predictors are equal to 0, β is a row vector with scalar Cox coefficients for each of the covariates, and x is a column vector representing covariate values. The covariates we used were the position of the previous reward, the port side (this nuisance variable was irrelevant for our interpretation of the data but nonetheless was necessary for accurate fitting), and photostimualtion condition (1 in stimulated trials and 0 otherwise).

We also used the fitted model parameters to simulate data. Similarly to the shuffled simulation described above, we first gathered all potential trials from the data. Next, to simulate leaving decisions we used the fitted Cox model to generate a sequence of probabilistic “coin-flips”, such that if one assumes “heads” to represent the hazard of leaving after poke n (taking into consideration its distance from the last reward, photstimulation condition etc.), then, for a given trial the simulation would go on with a series of coin-flips until the first “heads” is encountered - this would mark the time of leaving. In these simulations half of the data was used for fitting the model and the second half, for testing.

## Acknowledgements

We thank Bassam Atallah and Madalena Fonseca for comments on a previous version of the manuscript. We also thank Gil Costa for support with visual the diagram shown in Fig. 1. This work was supported by the European Research Council (Advanced Investigator Grants 250334 and 671251 to Z.F.M.), and Champalimaud Foundation (Z.F.M.).

## Author Contributions

E.L., D.B. and Z.F.M. designed the experiments. D.B., E.L., P.V., D.S. and M.O.L conducted the experiments. E.L. and P.V. analysed the data. E.L. and Z.F.M wrote the manuscript.

## Materials & Correspondence

Correspondence and requests for materials should be addressed to Z.F.M.

